# Use of linear features by red-legged partridges in an intensive agricultural landscape: implications for landscape management in farmland

**DOI:** 10.1101/2023.07.27.550774

**Authors:** Charlotte Perrot, Léo Séranne, Antoine Berceaux, Mathias Noel, Beatriz Arroyo, Léo Bacon

## Abstract

Current agricultural practices and change are the major cause of biodiversity loss. An important change associated with the intensification of agriculture in the last 50 years is the spatial homogenization of the landscape with substantial loss of such biodiversity-rich elements as seminatural linear features (hedgerows, field margins, grassy strips, etc.). In Europe, some management prescriptions serve to increase heterogeneity by the creation of these seminatural linear features which are not being used primarily for agricultural production. However, these elements are not equal in their support for biodiversity according to their structure and composition. The aim of this study is to determine the importance of landscape heterogeneity and specifically linear features on the spatial distribution of red-legged partridges, a small game species in decline in Europe. Through GPS-monitoring of adult birds, we first assess home range size throughout the year and during the breeding season, in relation to breeding status and to linear features (seminatural linear vegetation and tracks-roads for human traffic) density. Then, we focus on habitat selection during the breeding period in relation to linear features. We found that linear elements shape the use of space by red-legged partridges according to their reproductive status. Traffic routes and seminatural features structured by both herbaceous and woody cover, negatively influenced home range size. Further, breeding birds select linear elements with herbaceous cover while non-breeders select linear elements with woody cover, underlining the different needs of birds according to their breeding status. All birds selected areas near tracks, but non-breeders seemed to avoid roads. This study shows the importance for this species of the linear components that structure the agricultural landscape. We propose recommendations to promote the presence of the red-legged partridge in this agricultural environment but also of the biodiversity in general.

## Introduction

Agriculture is the most important land use activity, covering over a third of the world’s land surface (World Bank 2017). Current agricultural practices and changes are the biggest cause of biodiversity loss (Dudley and Alexander 2017) by having broad scale negative effects with the conversion of natural ecosystems into farms (Foley et al. 2005) and the release of pollutants (Brühl and Zaller 2019). An important change associated with the intensification of agriculture in the last 50 years is the spatial homogenization of the landscape with only a few sown crop types distributed in large uniform fields (Šálek et al. 2018) and with substantial loss of such biodiversity-rich elements as hedgerows, field margins and other non-cropped areas (Benton et al. 2003). For example, since 1950, 70% of the hedgerows have disappeared from the French *bocage* and continue to decrease. A fundamental concept in landscape ecology is that spatial homogeneity affects negatively ecological systems (Wiens 2002) which is also true for agricultural ecosystems.

In Europe, one response over biodiversity loss concerns in farmland has been the introduction of agri-environment schemes (AES), in which farmers are funded to modify their farming practices to provide environmental benefits (Batáry et al. 2015). Management prescriptions generally serve to increase heterogeneity (Benton et al. 2003) by the maintenance, creation or management of seminatural features which are not being used primarily for agricultural production. This includes for instance many linear elements like field margins, road verges, ditch banks, hedgerows, and wooded banks (Grashof-Bokdam and van Langevelde 2005). In intensively managed lowland landscapes, these linear features contain the greatest botanical diversity and provide different functions for wildlife. They act as corridors, facilitating species movement, providing shelters, food and nesting habitat for invertebrates (Lagerlöf et al. 1992; Delattre et al. 2013; Coulthard et al. 2016), reptiles (Knierim et al. 2018; Balouch et al. 2022), amphibians (Maes et al. 2008), mammals (Dondina et al. 2016; Pelletier-Guittier et al. 2020), and birds (Heath et al. 2017). The structure (height and width) and plant diversity of semi-natural linear features have also been shown to influence the biodiversity they support (Graham et al. 2018). Seminatural linear features are not equal in their support for biodiversity and this highlights the importance of guidelines for local stakeholders to manage them in line with their goals (Vickery et al. 2009; Montgomery et al. 2020).

The red-legged partridge (*Alectoris rufa*) is a typical species of agricultural land in southwestern Europe, where it has considerable socioeconomic value in rural environments as one of the principal small game species, including in France with ca 1.3 M birds hunted during the season 2013/14 (Aubry et al. 2016). However, despite its importance as a quarry species, populations have suffered marked declines during the last century throughout its range, including France (Souchay et al. 2022). Several reasons may explain these declines, but the most important appears to be habitat alteration, particularly changes occurring during recent decades in agrarian management systems (Delibes-Mateos et al. 2012). Previous studies on habitat preferences of partridges in agricultural environments have highlighted the importance of seminatural features for this species. For example, breeding density of red-legged partridge was found to correlate with the length of permanent field boundaries including hedgerows (Rands 1986; Meriggi et al. 1991) and with the amount of natural vegetation (Cabodevilla et al. 2021). Using radiotracking of breeding birds, Casas and Viñuela (2010) showed that herbaceous strips among fields were positively selected as nesting sites, with high nesting success. Today, to improve habitat selection studies, we can use new technological advances for higher spatial and temporal accuracy. Indeed, recent integration of high-resolution Global Positioning System (GPS) into miniaturized tracking tags has dramatically improved our ability to study habitat selection by providing us with accurate locations at high temporal frequency (Kays et al. 2015).

The aim of this study was to determine the importance of landscape heterogeneity and specifically linear features on the spatial distribution of red-legged partridges using GPS monitoring. Specifically, we first assessed home range size throughout the year and during the breeding season, in relation to breeding status and to linear features density. Then, we focused on habitat selection in relation to linear landscape features during the breeding period, and in relation to their structure: linear features considered included seminatural linear elements with either herbaceous or woody structure, and traffic routes since they are likely to influence wildlife habitat selection (Fahrig and Rytwinski 2009; Ascensão et al. 2012). We chose to consider distance from the linear feature into account, because not only the feature itself can be attractive or repellent to wildlife, but it can affect the suitability of nearby areas. For example, an area close to a hedge may be preferentially used, as a potential refuge is quickly accessible in the event of predation. Conversely, areas close to roads may be avoided because of the associated noise pollution. We expected breeding birds to be more selective than non-breeders as they are more constrained in their movements during the incubation and rearing period. Finally, we looked at how the nests of the tracked birds were located in relation to these linear features. Using the red-legged partridge as a flagship species, we discuss our results in terms of recommendations for landscape management in farmland areas to maximize benefits for breeding farmland birds.

## Methods

### Study area description and population

The study was conducted at two sites 56 km apart in Southwest France. The first is located in the village of Garganvillar in the Tarn-et-Garonne department (43°58’N, 1°30’E, 28.82 km2, hereafter TG Fig.1). The second is located within the villages of Azas, Montpitol and Roquesérière in the Haute-Garonne department (43°43’N, 1°39’E, 43.76 km2, hereafter HG, Fig.1). Both sites are characterized by an intensive agricultural landscape dominated by cereal crops (wheat, barley, oats), corn crops, leguminous crops (lentil, chickpeas) and oilseed crops (sunflower, rape). Their topography is relatively flat. These two sites benefit from a warm-temperate, oceanic climate with Mediterranean tendencies, characterized by a hot and fairly dry summer, a sunny autumn, a mild winter and a spring marked by recurring rainfall. Density of red-legged partridge is 2 pairs per 100 ha in TG and 1.5 pair per 100 ha in HG which is low. On both sites, captive-reared red-legged partridges are released for population reinforcement every year in July. Survival of released red-legged partridges to the following spring is lower than 0.05 (Souchay et al. 2018). Spring is also the period of capture for this study. Thus, we considered captured birds in spring as wild birds regardless of their origin. In the same way, because TG and HG have similar characteristics (topography, land use, climate and game management), and are relatively close, we did not differentiate the birds according to where they reside.

**Figure 1.**
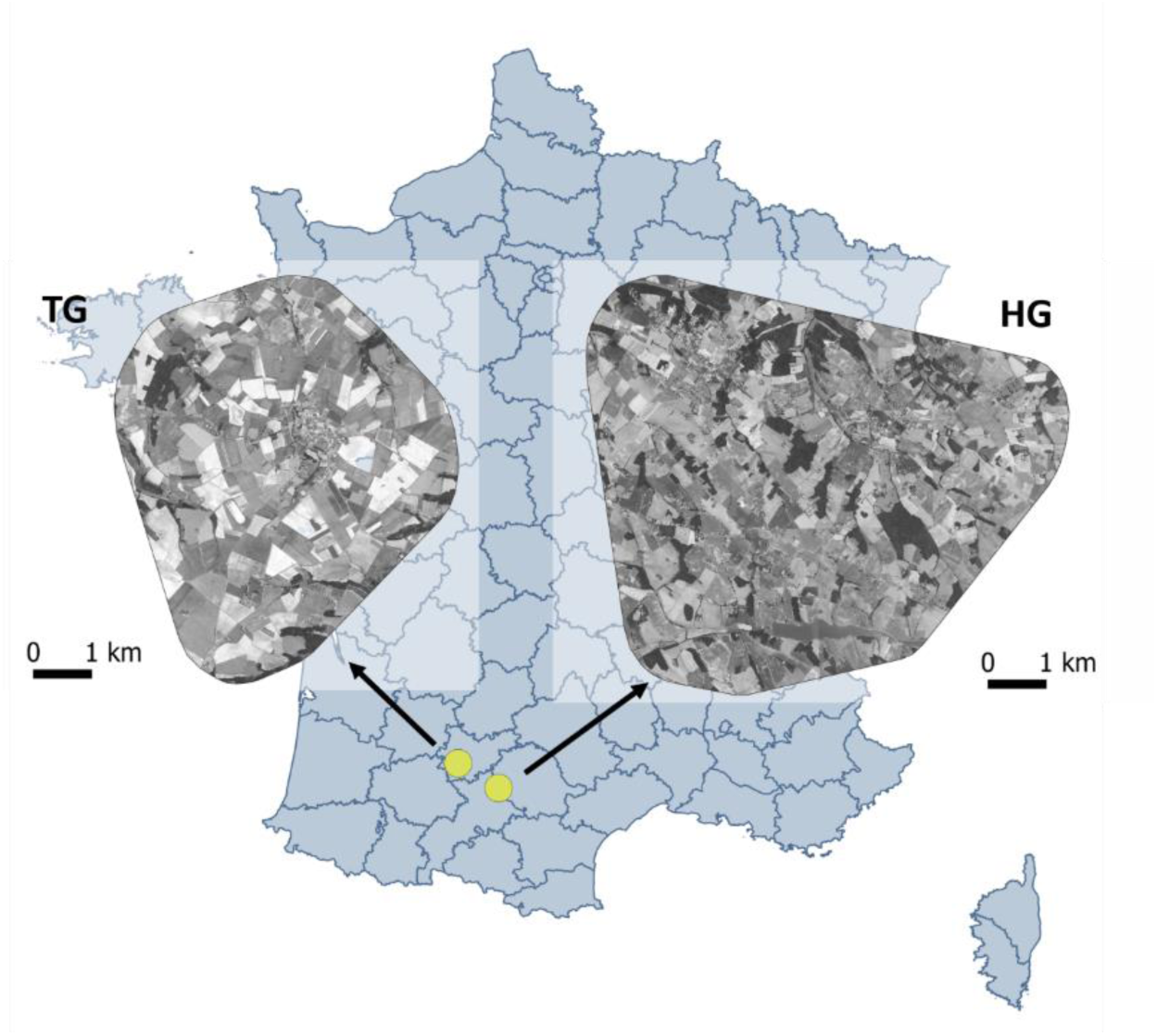
Map of the two study sites illustrating the intensive agricultural landscape.

The red-legged partridge exhibits a peculiar breeding system of double-nesting coupled with male incubation (Green 1984a; Casas et al. 2009). Females may lay eggs in two different nests, and when both clutches are laid, females usually start incubation of the second clutch while males incubate the first one and then raise their chicks (Green 1984a; Casas et al. 2009). Predators of the red-legged partridge are mostly generalist predators such as red foxes (*Vulpes vulpes*), feral cats (*Felis catus*), stone martens (*Martes foina*), free-roaming dogs (*Canis lupus familiaris*), and common buzzards (*Buteo buteo*) (Ferreras et al. 2022).

### Linear structure classification

Linear structures in the landscape include green linear features such as hedges and grassy strips but also human traffic routes. Green linear features were surveyed in the field from June to July 2018. They were classified into 4 categories according to the percentage of cover of the herbaceous stratum and the presence of a woody stratum. Herbaceous linear features were characterized by a herbaceous cover above 25% and by the absence of woody plants. Woody linear features were characterized by the presence of woody plants and by a herbaceous cover below 25%. Mixed linear features were characterized by an herbaceous cover above 25% and by the presence of woody plants. Finally, the last category was characterized by a herbaceous cover lower than 25% and by the absence of woody stratum. However, this category represented less than 1% of the vegetated linear feature, so was not considered in this analysis. Next, all green linear features were georeferenced using QGIS software.

Human traffic routes include roads (paved traffic routes) and tracks (unpaved traffic routes). To map roads and tracks in our two sites, we used vectors from IGN cartographic database (ROUTE 500® edition 2020) and from OpenStreetMap database (OSM). Then, for both sites and for all linear structures, distance-to-linear-feature rasters were created. Working with distances also allows to dilute the noise due to the imprecision of the GPS locations of the tracked birds, which is about 8 meters.

### Animal captures and GPS data

Wild red-legged partridges were caught using decoy traps (Smith et al. 1981). Traps are observed from about 100 meters so that the trapped partridge remained in the trap as little as possible. We equipped 16 birds in March and April 2018, and 17 birds in February, March and April 2019 with solar GPS-tags coupled to a UHF radio transmitters (e-obs digital telemetry, 10g). The device was attached to the bird with a field-adjustable leg-loop harness using Teflon ribbon. Birds were morphologically sexed. The origin of birds (wild vs. farmed) was also assessed by checking the condition of the plumage. Farmed birds show damaged plumage due to interactions with the wire mesh of their enclosure. However, this differentiation is only possible up to one year after their release, because once the breeding moult is done it is no longer possible to differentiate them. After release, birds were radio-tracked by triangulation using a portable Yagi antenna to download GPS data via the UHF system and monitor reproduction. Duration of monitoring was very variable between individuals (mean ± SD = 111 ± 159 days), due to the natural mortality of birds, UHF signal loss and GPS device dysfunctioning. All the animals were treated according to the ethical conditions detailed in the specific accreditations delivered to the Office Français de la Biodiversité by the Préfecture de Paris (prefectorial decree n°2009-014) in agreement with the French environmental code (Art. R421-15 to 421-31 and R422-92 to 422-94-1).

From March to October, GPS devices recorded locations every hour from 3:00 to 21:00 UTC, that is 19 locations per day. From November to February, as days are shorter, we lightened the programming to avoid draining the battery. During this time, GPS devices recorded locations every hour from 6h00 to 18h00 UTC, that is 13 locations per day. For all GPS data, we screened them for erroneous locations from the individuals’ trajectories (Bjørneraas et al. 2010). To avoid noise from the capture and death of the bird, we removed the first and last day of monitoring.

Reproductive status of partridges was estimated by the detection of incubation. An incubating bird was considered as breeding and conversely a non-incubating bird was considered as non-breeding. Detection of incubation was done by two methods. First, in 2018, birds were radio-tracked and thus observed twice a week to detect if they are incubating or not. When a bird was located 3 times successively in the same place in the field, it was considered in incubation and an estimation of the nest location was recorded. Once incubation was complete, the field agent checked for the presence of the nest. In 2019, for lack of human resources, field observations could not be done. We therefore identified nesting attempts and nest location from bird trajectories using the *find_nests* function of the *nestR* package (Picardi et al. 2020). We have previously verified the robustness of this method by comparing the nest locations of birds equipped in 2018 estimated by this method with the locations recorded for these same nests in the field.

### Analysis

#### Annual space use

For monthly home range size estimation, we selected individuals tracked at least one full month, that is 17 individuals (8 females with 4 breeders and 4 non-breeders, and 9 males with 3 breeders and 6 non-breeders). Mean monitoring length of these birds was 4.9 (± 4.2 SD) months. We derived home range size for each individual for each month from an utilization distribution (UD) computed using the Brownian bridge movement model (Horne et al. 2007). BBMM is a continuous time stochastic model of movement that incorporates an animal’s movement path and time between locations to calculate UD. We calculated monthly UDs at 95% using the *R adehabitat* package (Calenge 2011). For each home range, the density of each linear structure (roads, tracks and green linear features) was calculated in meters of linear structure per hectare using QGIS software (version 3.10.4-A Coruña).

Next, we investigated if the month of the year, the individual characteristics and the density of linear structures had an influence on the variability of monthly home range size using linear models. Since we had a relatively small sample size and since males are likely to incubate and participate in the rearing of young, we chose to consider only the reproductive status of the individual and not their sex. In other words, if a male or female individual was seen incubating, it was considered a breeder, if not as a non-breeder. In the same way, as we did not know the origin (wild or farmed bird) for all individuals, we could not test its influence on the monthly home ranges size. Therefore, the complete model contained the month of the year, the reproductive status of individual and density of the five linear structures as explanatory variables. Since there may be multiple monthly home ranges per individual, bird identity was included as a random effect. Next, we tested the significance of this random effect by the function *ranova* of the R package *lmerTes*t (Kuznetsova et al. 2017) to decide on the relevance of maintaining this effect in the model. To satisfy the assumptions of the model (i.e. normality and homoscedasticity of residuals), home range size was log-transformed. Following recent recommendations to produce model estimates comparable between and within studies, we standardized all explanatory variables by centring and dividing by two standard deviations. To check for overparameterization, we used the cross-validation method from the *caret* R package (Kuhn 2005). Model selection was based on Akaike Information Criterion corrected for small sample size (AICc). When several models were within a Δ AICc of 2 from the best model, we employed a model averaging approach, using the natural average method implemented in the *MuMIn* package (Grueber et al. 2011) of R on models within two points of AICc from the best one. This allowed us to account for model selection uncertainty in order to obtain robust parameter estimates.

#### Space use and habitat selection during the breeding period

To study space use during the breeding period we focused on home range size, nest location and habitat selection during breeding. We chose the period when most individuals were monitored simultaneously and over the same duration in 2018 and 2019. This period ranges from March 26 to June 5 in 2018 and 2019, therefore 71 days of monitoring for 10 tracked individuals (4 females all breeding and 6 males with 2 breeders et 4 non-breeders). It includes the end of the pair formation, nest construction and incubation.

#### Home range

We calculated home range size for every individual during this period using the same method as for monthly home range size. As before, we investigated if the month of the year, individual characteristics and the density of linear structures had an influence on the variability of this home range size. First, we tested our hypothesis that non-breeding birds should express a larger home range than breeding birds. Hence, we compared these two groups using the non-parametric Mann-Withney-Wilcoxon test using the alternative “greater.” Next, we tested the influence of month and green linear features on home range size using linear models. Since the sample size is small (N=10), we tested each variable one by one, which makes six different models. To satisfy the assumptions of the model (i.e. normality and homoscedasticity of residuals), home range size was log-transformed.

#### Nest habitat

Distance from each nest to each linear structure was calculated. When distance between the nest and the green linear feature was less than 8 meters (GPS accuracy), we considered that the nest was within this linear feature. Then we checked if the nest was close to a road or a track.

#### Habitat selection

Since red-legged partridges are mainly active during the day, and since we had few night locations, we used only the locations recorded during daytime. To evaluate how linear structures influence habitat selection, observed 60-min movement steps (i.e. straight lines connecting successive GPS locations) were compared to random steps; the former are typically used to estimate habitat use, and the latter as an estimate of habitat availability (Thurfjell et al. 2014). Habitat selection and movement are interlinked, with movement rates influencing selection and vice versa (Forester et al. 2009). Therefore, integrated step-selection analysis was used to compare observed and random steps to evaluate habitat selection while simultaneously accounting for the differing movement behaviours (Avgar et al. 2016).

Twenty-five random steps for each observed step were generated using analytical distributions that were parameterized based on the observed animal movement patterns. Observed movement patterns were described using gamma distribution of step length and a Von Mises distribution of turning angles from each individual (Signer et al. 2019).

Habitat selection was evaluated as a function of distances to linear structures: green linear features (herbaceous, woody, mixed), roads and tracks. Habitat attributes at the end of each observed step were compared to attributes at the end of random steps. Distances were transformed using the natural logarithm because we expected animals to respond more strongly to features closer to them, with the response declining at an unknown rate (Dickie et al. 2020). Each individual was modelled separately using conditional logistic regression (*amt* package in R ; Signer et al. 2019). The relative probability of selection was modelled as a function of distance linear structures. The step length, natural logarithm of step length, cosine of turning angle were included as modifiers of the observed movement parameter.

Individual selection and movement responses were summarized to evaluate consistency in responses based on model coefficients and their 95% confidence intervals (CIs). Selection was defined as occurring if use was higher than availability, resulting in positive selection coefficients, and avoidance if use was less than availability, resulting in negative selection coefficients. CIs overlapping with zero were interpreted as indifference, and non-overlapping CIs as significant selection/avoidance depending on the variable of interest (Dickie et al. 2020; Fieberg et al. 2021).

Then, to test the influence of breeding status on habitat selection, we used inverse variance-weighted linear modelling to obtain population-level averages for both groups (Murtaugh, 2007). Finally, to understand how strongly linear features influenced selection, the relative selection strength was calculated (Avgar et al. 2017). For all linear structures, we calculated the relative probability of selecting an area according to its distance with these structures.

## Results

### Linear structure length

On both sites, 147.25 km of roads and 63.91 km of tracks were recorded. Concerning green linear features, 52.39 km, 11,18 km and 65.68 km were respectively recorded for herbaceous, woody and mixed linear features.

### Annual space use

The mean size of monthly home range was 25.14 ha ± 1.28 SE. Four models explained home range size variation almost equally well (Table 1). Model averaging indicated a negative effect of the density of the two traffic ways, although this effect was less marked for roads than for paths (Table 2, Fig 2a, 2b). In the same way, density of mixed linear features negatively influenced home range size (Table 2, Fig 2c). We did not detect any effect of reproductive status, month and density of herbaceous and woody linear feature densities on monthly home range size.

**Figure 2.**
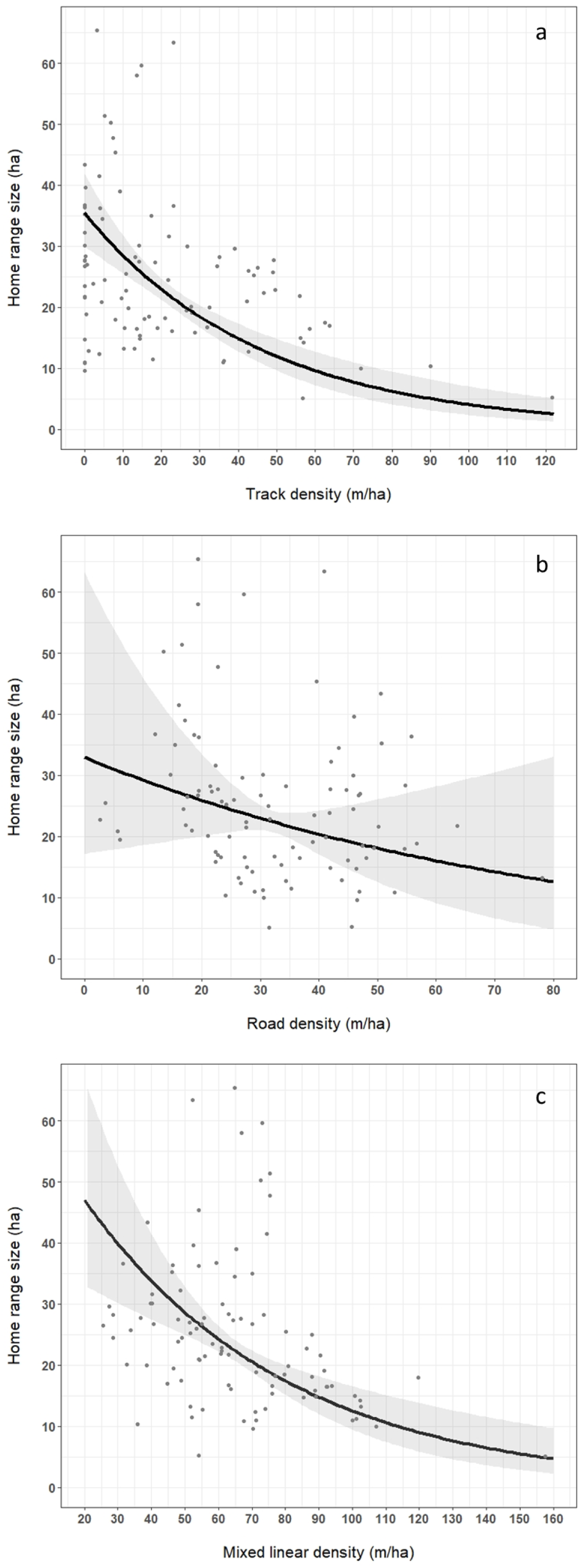
Linear relationship between home range size and a) track density, b) road density, c) mixed linear feature density on monthly in red-legged partridge according to the model averaging of the top model set (estimate ± 95%IC). In grey, points represent raw data.

**Table 1.** Model selection of the factors influencing monthly home range size in red legged partridge. Models with a Δ AICc ≤ 2 from the best model are represented. LF = linear feature.

| Response variable | Models | df | logLik | AICc | delta | weight |
| --- | --- | --- | --- | --- | --- | --- |
| Home range size (log) |  |  |  |  |  |  |
|  | *track density + road density + mixed LF density* | 5 | -41.14 | 92.95 | 0 | 0.37 |
|  | *track density + HW density + herbaceous LF density* | 5 | -41.35 | 93.38 | 0.43 | 0.30 |
|  | *track density + road density + mixed LF density + herbaceous LF density* | 6 | -40.73 | 94.41 | 1.46 | 0.18 |
|  | *track density + road density + mixed LF density + woody LF density* | 6 | -40.82 | 94.60 | 1.65 | 0.16 |

**Table 2.** Model-averaged estimates ± SE and 95%CI of parameters explaining variations in monthly home range size in red-legged partridge. The relative importance of each factor is calculated by summing the AIC weights across the top models (Table 1) where the given factor appears (last column). LF = linear feature.

| Response variable | Parameters | Estimate | SE | Confidence interval | Sum of weight |
| --- | --- | --- | --- | --- | --- |
| Home range size (log) |  |  |  |  |  |
|  | **Track density** | **-0.50** | **0.08** | **(-0.66 ; -0.33)** | **1** |
|  | **Road density** | **-0.24** | **0.11** | **(-0.47 ; -0.02)** | **0.70** |
|  | **Mixed LF density** | **-0.37** | **0.09** | **(-0.55 ; -0.20)** | **1** |
|  | *Herbaceous LF density* | -0.21 | 0.13 | (-0.46 ; 0.04) | 0.47 |
|  | *Woody LF density* | -0.06 | 0.08 | (-0.22 ; 0.10) | 0.16 |

### Reproductive space use

#### Home range

While not statistically significant at the 0.05 level, the Mann-Withney-Wilcoxon test indicated that the home range size of non-breeders tends to be larger than that of breeders (W = 20, p = 0.057). During the reproductive period ranging from March 26 to June 5, non-breeders had a mean home range size of 53.19 ha ± 12.60 SE while breeders had a mean home range size of 22.34 ha ± 3.27 SE (Fig. 3). Concerning linear structures, track density and mixed linear feature density negatively influenced home range size (tracks: F_18_ = 7.33, p = 0.03; mixed linear features: F_18_ = 27.51, p = 0.0008, Fig. 4a, 4b). We did not detect any effect of road, woody and herbaceous linear feature density on reproductive home range size.

**Figure 3.**
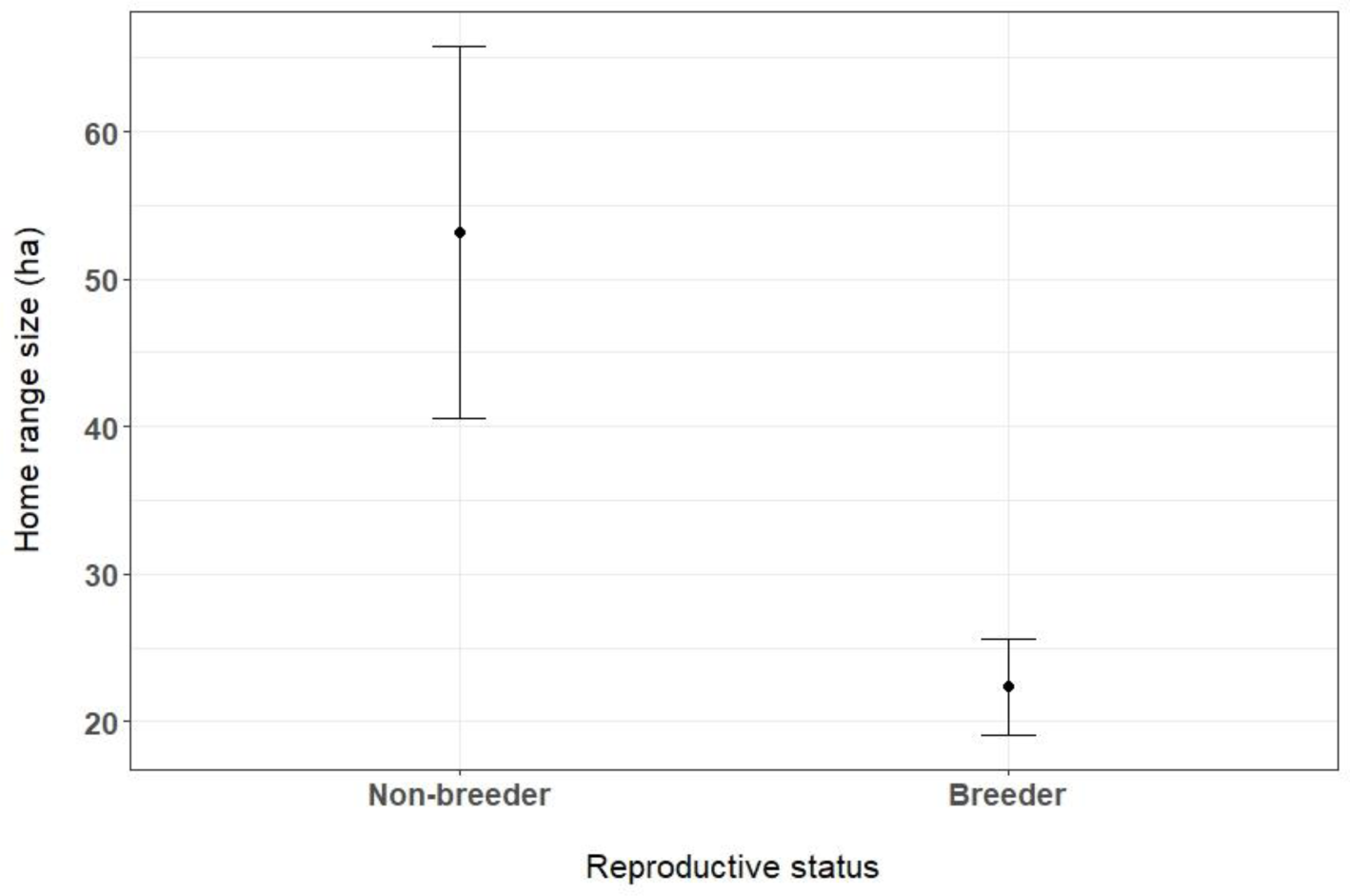
Mean home range size ± SE of red-legged partridge for non-breeders (N=4) and breeders (N=6).

**Figure 4.**
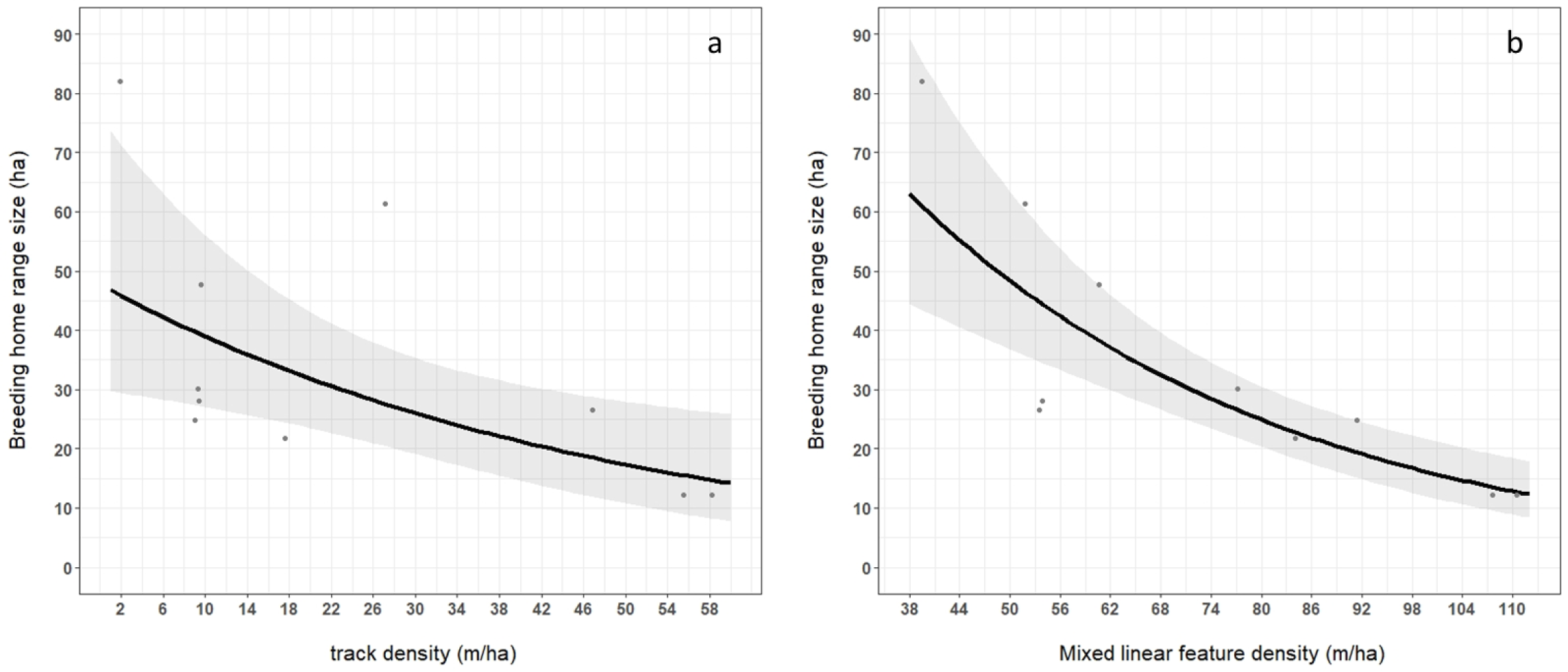
Effect of a) track density, b) density of mixed linear feature on home range size during breeding in red-legged partridge according to linear regression models (estimate ± 95%IC). In grey, points represent raw data.

#### Nest habitat

9 nest locations of 7 different breeders were identified: 7 by field observations and 2 more in 2019 by studying the trajectory of the birds using the *nestR* package. The low number of nests found using *nestR* package in 2019 is probably due to the difficulty of determining incubation attempts when these quickly fail. 4 of these nests were located in mixed linear features, of which 2 were close to a road (distance = 6.26 m and 7.14 m) and 2 close to a track (distance = 1.18 m and 6.02 m). 4 other nests were located in herbaceous linear features, all near a road (distance = 3.72 m, 3.20 m, 4.47 m and 7.71 m). The last nest was located in a garden containing grasses and bushes, but no linear structures.

#### Linear features selection

Breeding red-legged partridges selected to be closer to herbaceous and mixed linear features and to tracks. However, they did not show any selection or avoidance behaviours concerning woody linear features and roads (Table 3). Concerning distance to herbaceous linear features, it was 1.90 times more likely that an individual would use areas 5 m away from this linear feature than an area 50 m away, which itself was 1.47 times more attractive than an area 200 m away. For distance to mixed linear features, it was 1.68 times more likely that an individual would use an area 5 m away from this linear than an area 50 m away, which itself was 1.37 times more attractive than an area 200 m away (Fig. 5a). In the same way, the probability of using an area 5 meters from a track was 1.25 times higher than that of using an area 50 meters from the track, which itself was 1.14 times more attractive than an area 200 meters away (Fig. 5a).

**Figure 5.**
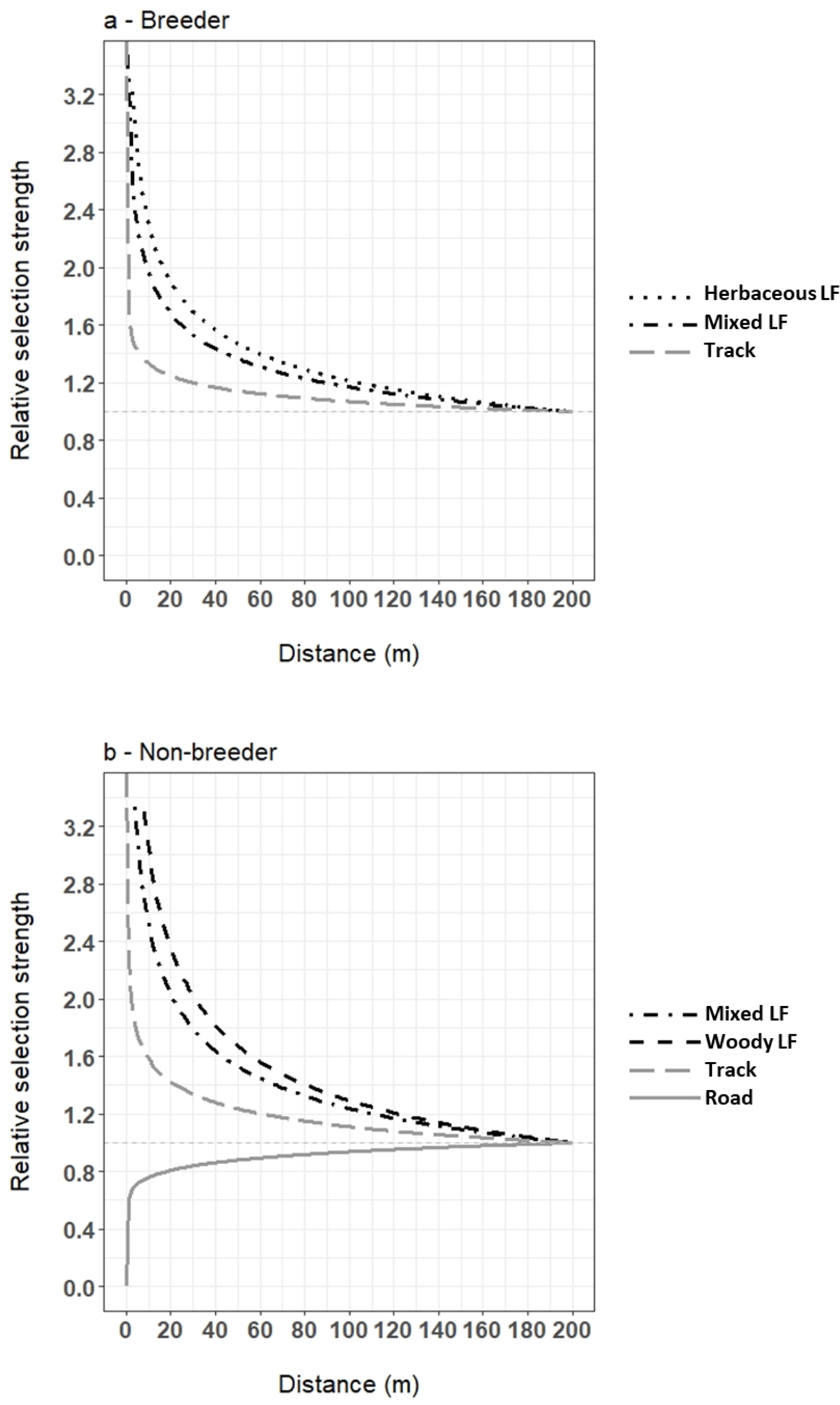
Relative selection strength of herbaceous (dotted line), mixed (dotted-dashed line), woody (dashed line) linear features and of tracks (grey long dashed line) and roads (grey solid line) for breeder (a) and non-breeder (b) red-legged partridges.

**Table 3.**
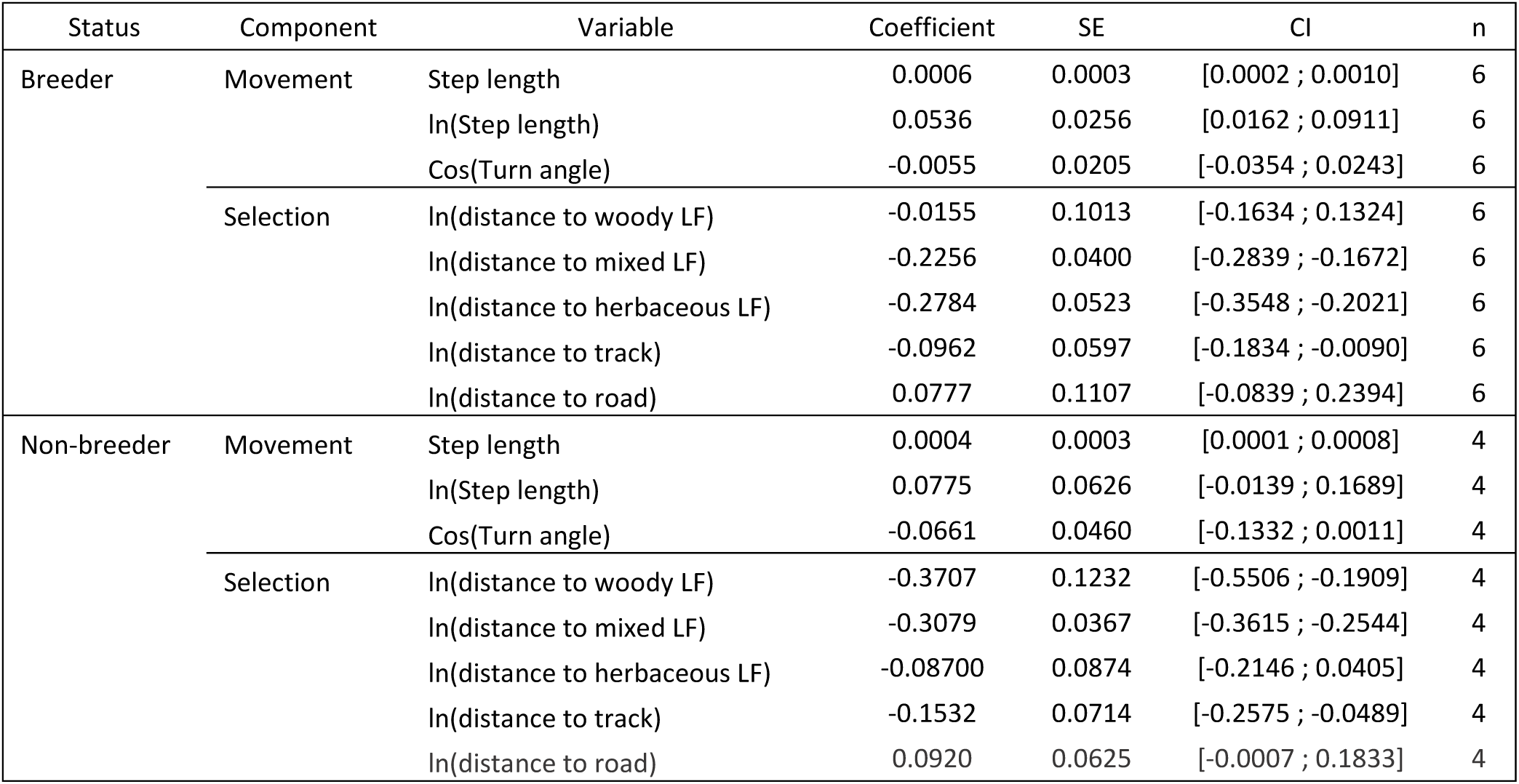
Model-averaged estimates ± SE and 95%CI of parameters explaining variations in monthly home range size in red-legged partridge. The relative importance of each factor is calculated by summing the AIC weights across the top models (Table 1) where the given factor appears (last column). LF = linear feature.

Non-breeder birds selected to be closer to woody and mixed linear features and to tracks but they were indifferent to herbaceous linear features. (Table 3). They also tended to avoid the areas closest to roads (Fig. 5b). An area 5 meters away from a woody linear feature was 2.35 times more attractive than an area located 50 meters away, which was itself 1.67 times more attractive than an area located 200 meters away (Fig. 5b). For mixed linear features, an area 5 meters away from was 2.03 times more attractive than an area located 50 meters away, which was itself 1.53 times more attractive than an area located 200 meters away (Fig. 5b). In the same way, the probability of using an area 5 meters from a track was 1.42 times higher than that of using an area 50 meters from the track, which itself was 1.24 times more attractive than an area 200 meters away (Fig. 5b).

## Discussion

The reproductive status of birds, as well as the structure of the habitats in which they live, significantly influence their use of space (Rolando 2002). Our study is consistent with this, as tracked red-legged partridges occupied space differently depending on their reproductive status. First, during the breeding season, breeding birds tended to have a smaller home range than non-breeders, probably due to the constraints of breeding. Indeed, during the incubation period, the movements of the nesting males and females are limited to the nest and its close environment. Then, during the rearing period, parents must adapt to the low movement abilities of their non-flying chicks (Green 1984b).

Second, reproductive status also shapes habitat selection. In spring, semi-natural linear features with herbaceous cover were highly selected by breeders. Moreover, all monitored nests but one were located in this type of linear feature, i.e. those with an herbaceous stratum. These results are consistent with previous studies on partridges that have shown the importance of grass cover for nesting (Rands 1986; Casas and Viñuela 2010). Indeed, field margins with herbaceous plants such as grassy strips, provide a cover to hide the nest from predators, food for adults (grasses and seeds) and for chicks (arthropods) (Vickery et al. 2009). However, field margins are known to attract predators because of their foraging behaviours toward linear features (Morris and Gilroy 2008). For example, during foraging, field margins and hedgerows facilitate movement of foxes, a partridge predator, through the landscape (Feber et al. 2019). Hence, nesting in these linear landscape elements could be an ecological trap like for the grey partridge (Bro et al. 2004; Rantanen et al. 2010). Conversely, non-breeding birds chose linear elements with a woody structure (bushes and/or trees) with or without grass cover. Non-breeders chose these hedgerows probably because the woody structures provide cover against raptors and they can exploit nearby resources or access to locations which might otherwise be too risky to use at all (Hinsley and Bellamy 2000). Hedgerows with a woody and herbaceous stratum were therefore selected by red-legged partridges regardless of their reproductive status. Furthermore, we found in our tracked partridges that the higher the density of these hedges, the more the monthly and breeding home range were reduced. These two results together underline the importance of these hedges with woody and herbaceous strata for the red-legged partridge, probably for its multiple functions (shelter, food resource, movement facilitator). Moreover, increasing availability of these semi-linear elements could also dilute predation pressure on those elements.

In the same way, both breeders and non-breeders selected tracks or areas very close to them, and home range size was negatively correlated with track density. There are several non-exclusive hypotheses for the attraction of unpaved roads. Firstly, tracks may be unobstructed ways that facilitate the movement of ground-dwelling birds such as partridges. Secondly, tracks may also be associated with grassy ditches and banks, or hedgerows. Thus, partridges potentially find shelter and food closer to tracks. Finally, tracks are a source of grit used by partridges mainly for grinding hard food items but also for its nutritional role as a source of minerals (Gionfriddo and Best 1999).

Compared to tracks, roads have more intense traffic which is a source of disturbance due to the noise, but also a source of mortality (Forman and Alexander 1998). Hence, in our study, non-breeding red-legged partridges tended to avoid these noisy and hazardous habitats probably to ensure their survival. This result is consistent with the study of Cooke et al. (2020) that found a negative relationship between red-legged partridge abundance and road exposure. Moreover, the regular discovery of red-legged partridges killed on the roads by frequent roadkill events in our study site, gives weight to this hypothesis (unpublished data). However, breeding red-legged partridges did not avoid the roads contrary to non-breeding red-legged partridges. In addition, four of the eight nests found were located within 8 meters of a road. Thus, the road environment appears to attract breeding individuals during reproduction. Similar to tracks, roads are frequently surrounded by ditches and grassy banks full of seeds and invertebrates, providing nesting habitat as well as food for adults and chicks. However, many studies demonstrate that road-side productivity and plant diversity attract wildlife while at the same time inflicting mortality so that roads can act as ecological traps (Coffin 2007). Hence, the use of this sub-optimal habitat by the red-legged partridge is probably due to the scarcity of favorable nesting sites in this context of intensive agriculture. Surprisingly, the partridges with a small home range were the ones with the highest road density in their home range, although this relationship is tenuous. Thus, we could think that the road environment provides the necessary resources for red-legged partridges during the year. But it is more likely that roads act as dangerous barriers that impede movement and thus limit the extent of bird territories. Red-legged partridges seem affected by the habitat fragmentation through the development of the road network like in other terrestrial bird species (Laurance et al. 2004).

Red-legged partridges seem also impacted by the homogenization of agricultural landscapes due to the development of intensive agriculture. This homogenization results into larger fields, and thus rarer semi-natural lines (hedgerows, field sides and unpaved roadsides). Red-legged partridges must move further to cover their basic needs (food, nest, resting place, etc.) leading to an increase in their home range size as in other farmland species (Michel et al. 2017; Mayer et al. 2022). This could explain why home ranges estimated in our study are up to 2.7 times larger than those previous studies during the breeding season (Ricci 1985; Mauvy et al. 1991). But monitoring methods may also be the cause of this size discrepancy. GPS tracking allows the recording of a greater number of locations than the radio tracking used in previous studies, which may explain the larger home range in our study.

Finally, we found no seasonal effect on monthly home range size, which is surprising because (1) birds generally have different movement constraints in relation to their phenology (breeding, dispersal, fall company) (2) the seasonal rotation of agricultural crops changes the distribution of food resources for farmland species during the year. Due to the small number of birds monitored, the lack of power in our analyses could explain the lack of detection of this seasonal variability. Moreover, our small sample size does not allow us to control for potential study site nor year effects. It is important to take this limit into account when interpreting the results, particularly those concerning habitat selection during the breeding period. Finally, in addition to increasing the sample size, better characterizing these linear elements by taking into account their size (height, length, width), their specific plant composition or their age, would enable more precise and effective recommendations to be made.

### Management recommendations

Orłowski (2008) proposed to move bushy vegetation away from the roads to reduce the likelihood that birds would cross the road and die. In the same way, we recommend the creation of a vegetation-free buffer zones along roadsides to prevent ground-nesting birds from colliding with vehicles but in roadkill hotspots only. Vegetation removal in roadkill hotspots should always be combined with small-mesh fencing to prevent passage of not only red-legged partridges, but also their predators (foxes) and small species mammals that inhabit the roadside (Galantinho et al. 2022). However, on roads with less risk of collision, this roadside management does not seem appropriate because it would be to the detriment of many species, since vegetated roadsides are favourable to the establishment of small mammals (Ascensão et al. 2012), insects (New et al. 2021) and contribute to the plant diversity (Lázaro-Lobo and Ervin 2019). Hence, collision risk mapping in this intensive agricultural environment would greatly facilitate roadside management.

For the red-legged partridge we believe that the major problem is not road layout but the lack of semi-natural elements in these intensive agricultural landscapes, so that birds are forced to use non-optimal habitats such as roadsides. These results underline the importance of developing a network of semi-natural elements in agricultural areas, such as hedges, grassy strips or embankments for the preservation of the red-legged partridge and of biodiversity in general. We have seen that, depending on their reproductive status, red-legged partridges did not select the same kind of field margins because they do not provide the same resources (nesting site, food, shelter, corridor, etc.). More broadly, the quantity of seminatural elements and their heterogeneity in shape, size, structural complexity, and also in their plant composition are essential to meet the needs of the different species living in this agricultural environment. For example, hedgerows favoured the presence of forest edge birds (Mallet et al. 2022) but may also be detrimental for some grassland bird species, such as skylark (*Alauda arvensis*) for which grass strips are beneficial (Josefsson et al. 2013). Equally, some bats are more active in hedgerows composed mainly of trees (Lacoeuilhe et al. 2018), while the hazel dormouse (*Muscardinus avellanarius)* and the European badger (*Meles meles*) prefer hedgerows with a high shrub cover (Dondina et al. 2016).

Agricultural policies should therefore support the increase of linear semi-natural areas in agricultural landscapes without forgetting to consider their complexity and heterogeneity so has to better meet management and conservation objectives. The maintenance over time of these seminatural linear elements should be part of this policy, so that they continue to fulfil their function for biodiversity. For this to work, it is essential to get farmers on board with this biodiversity conservation effort. So, it is necessary to measure the effectiveness of this habitat management, by monitoring the fauna and flora and to transfer these results from scientists to local stakeholders.

## Acknowledgements

We would like to thank all the people involved in the fieldwork among sites and over years and in particular Frédéric Le Capitaine, Benoît Laboup, Arnaud Gaujard, Julie Grezeleau, Romain Mongeot, Estella Vignon, Anne-Charlotte Pommier-Petit, Soumaya Belghali and Lisa Gili. We would also like to thank our partners for their support in this study: the Federation of Hunters of Haute-Garonne (FDC 31) and the Federation of Hunters of Tarn-et-Garonne (FDC 82). We thank Aude Géraud for her logistic support during the project. We also would like to thank Françoise Ponce, Jean-Bernard Puchala and Luc Fruitet for their help in setting up this project. We are grateful to Michel Ruas for the donation of 2 farmed red-legged partridges which allowed us to improve our skills in equipping birds with GPS tags.

## Data, scripts, code, and supplementary information availability

Data, scripts and code are available online: https://doi.org/10.5281/zenodo.8144130

## Conflict of interest disclosure

The authors declare that they comply with the PCI rule of having no financial conflicts of interest in relation to the content of the article.

## Funding

This project was funded by the OFB, Agrifaune and the Regional Federation of Hunters of Occitanie.

## Notes

https://doi.org/10.5281/zenodo.8144130

### Competing Interest Statement

The authors have declared no competing interest.

### Summary of Updates

The PCI Ecology badge was added on the 1st page of the article. It indicates that the article has been peer-reviewed and recommended by PCIEcology.

